## Supplemental Information for "Convergent eusocial evolution is based on a shared reproductive groundplan plus lineage-specific plastic genes"

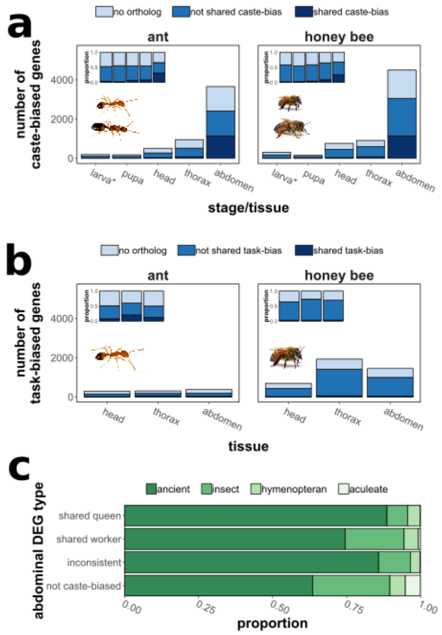


**Supplementary Figure 1. Patterns of caste-biased expression in pharaoh ants and honey bees.**

Deleted: Supplementary Fig.

Number of differentially expressed genes (FDR < 0.05) between a) queens and workers and b) nurses and foragers at each developmental stage or tissue in ants (left) and honey bees (right). “Head”, “thorax”, and “abdomen” refer to body segments of adults, while “pupa” and “larva” refer to whole bodies. “No ortholog” refers to genes for which no 1:1 ortholog exists (either due to apparent duplication or complete lack or orthology), “not shared caste/task bias” refers to genes for which 1:1 orthologs can be identified but are only differentially expressed in one species, and “shared caste/task” bias refers to genes for which 1:1 orthologs are differentially expressed in both species. Insets show the proportion of each category of gene out of all differentially expressed genes at that stage or tissue. c) Proportion of abdominal DEGs by estimated evolutionary age (shading). “Shared queen/worker” indicates genes upregulated in queen or workers of both species. Source data are provided as a Source Data File, “Supplementary_Fig1a.txt”, “Supplementary_Fig1b.txt”, “Supplementary_Fig1c.txt”.
*****: the category “larva” represents differential expression across larvae of all stages for which caste can be identified (second to fifth larval stage). Photos were taken by Luigi Pontieri (pharaoh ants) and Alex Wild (honey bees).

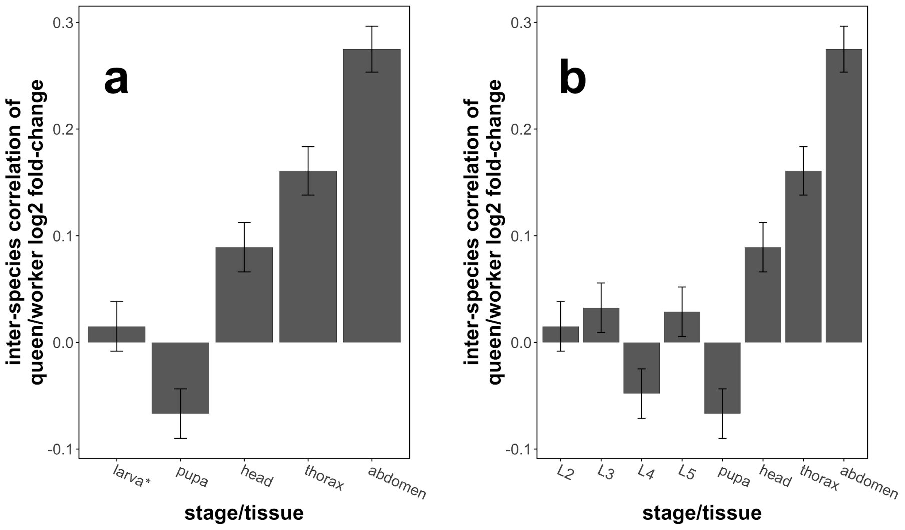


**Supplementary Figure 2. Correlation of gene-wise $log_2$ fold-change between queens and workers in ants and honey bees.**
Pearson correlation of $log_2$ fold-change between queens and workers as measured at each stage or tissue in *M. pharaonis* and *A. mellifera* for each 1:1 ortholog (N = 7640). Error bars indicate Pearson correlation 95% confidence intervals. In (a), the category "larva*" represents differential expression across larval stages, while in (b) each larval stage (L2-L5) is plotted individually. Source data are provided as a Source Data File, "Supplementary_Fig2a.txt", "Supplementary_Fig2b.txt".

**Deleted: Supplementary Fig.**

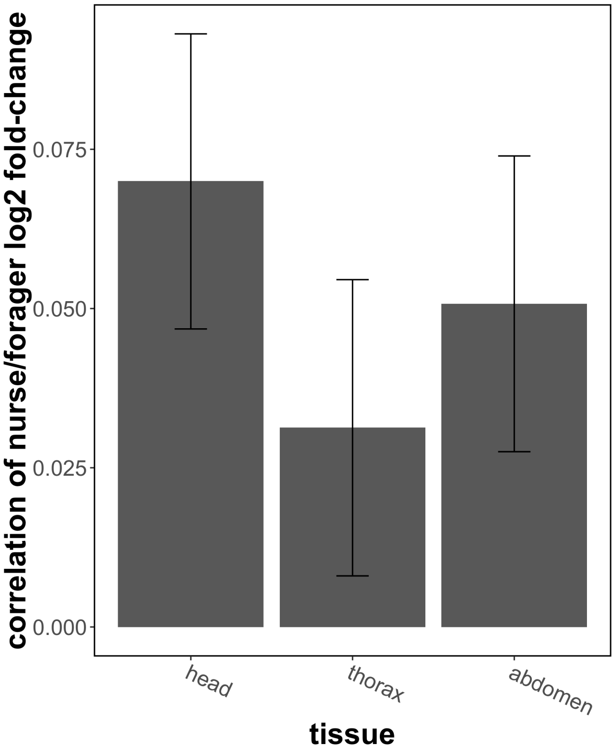


**Supplementary Figure 3. Correlation of gene-wise log$_2$ fold-change between nurses and foragers in ants and honey bees.**

Deleted: Supplementary Fig.

Pearson correlation of log-fold change between nurses and foragers as measured in each tissue in *M. pharaonis* and *A. mellifera* for each 1:1 ortholog (N = 7640). Error bars indicate Pearson correlation 95% confidence intervals. Source data are provided as a Source Data File, “Supplementary_Fig3.txt.

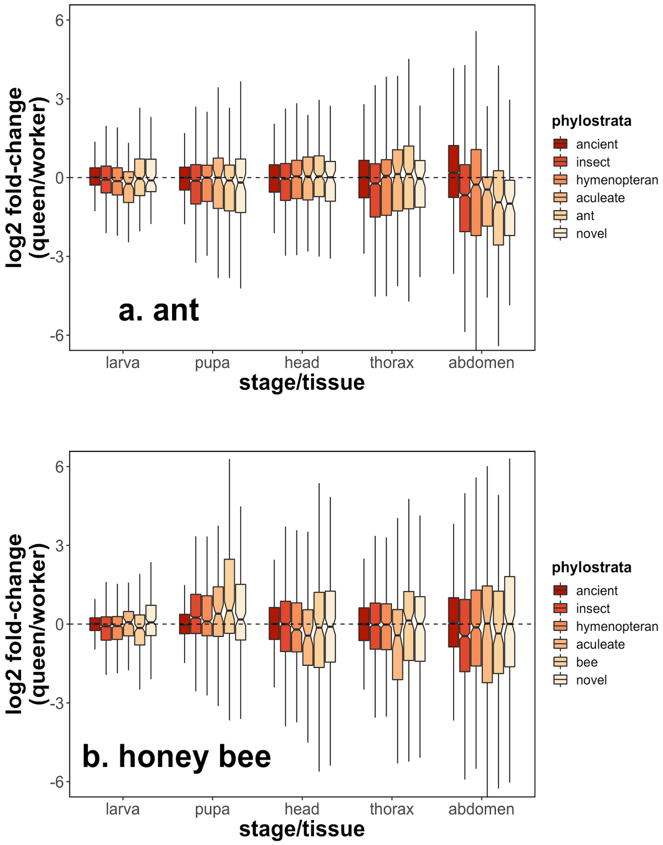


Supplementary Figure **4. $Log_2$ fold-change between queens and workers across phylostrata.** $Log_2$ fold-change at each stage/tissue in each phylostrata category. Positive values indicate higher expression in queens compared to workers. $Log_2$ fold-change has been adjusted relative to the median value at that stage/tissue, in order to compare across tests. "Ancient" genes indicate any genes shared beyond insects (i.e. with vertebrates). $Log_2$ fold-change varies according to phylostrata for every stage/tissue for each species (LM; LRT; $P < 0.001$). In each boxplot, the middle line represents median values, outer edges of boxplot represent upper and lower quartiles, and whiskers represent a deviation of 1.5*(interquartile range) from the upper and lower quartiles. Source data are provided as a Source Data File, "Supplementary_Fig4a.txt", "Supplementary_Fig4b.txt".

**Deleted: Supplementary Fig.**

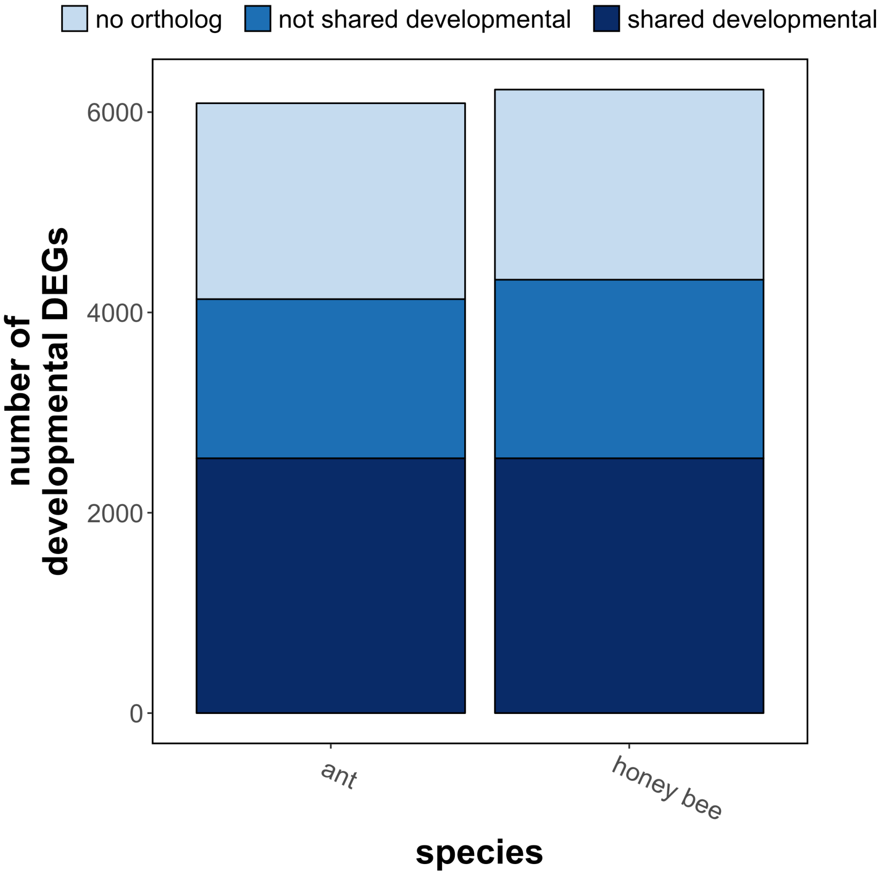


**Supplementary Figure 5. Overlap of developmentally-biased genes in ants and honey bees.** Number of developmental differentially expressed genes in ants and in honey bees. “No ortholog” refers to genes for which no 1:1 ortholog exists (either due to apparent duplication or complete lack or orthology), “not shared developmental” refers to genes for which 1:1 orthologs can be identified but are only differentially expressed between developmental stages in one species, and “shared developmental” refers to genes for which 1:1 orthologs are differentially expressed between embryonic and larval developmental stages in both species. Source data are provided as a Source Data File, “Supplementary_Fig5.txt”.

**Deleted: Supplementary Fig.**

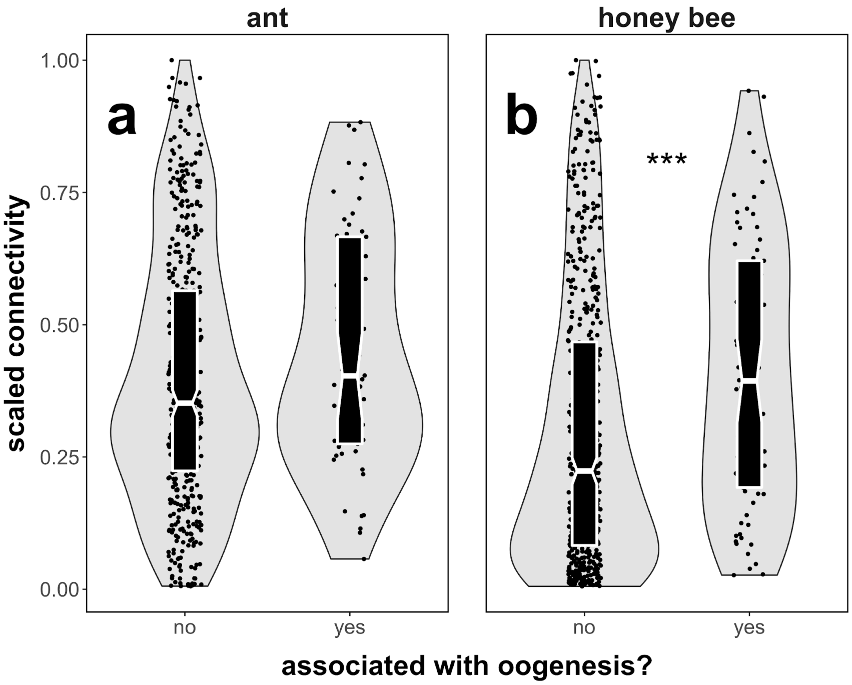


**Supplementary Figure 6. Genes associated with oogenesisis are more highly connected within queen abdominal modules.**

Deleted: Supplementary Fig.

Orthologs of genes associated with oogenesis in *D. melanogaster* are more highly connected within queen abdominal modules in honey bees (*** = P < 0.001; Wilcoxon test) though not in ants (P = 0.114). N = 542 (ants), 649 (honey bees). ). In each boxplot, the middle line represents median values, outer edges of boxplot represent upper and lower quartiles, and whiskers represent a deviation of 1.5*(interquartile range) from the upper and lower quartiles. Source data are provided as a Source Data File, “Supplementary_Fig6.txt”.

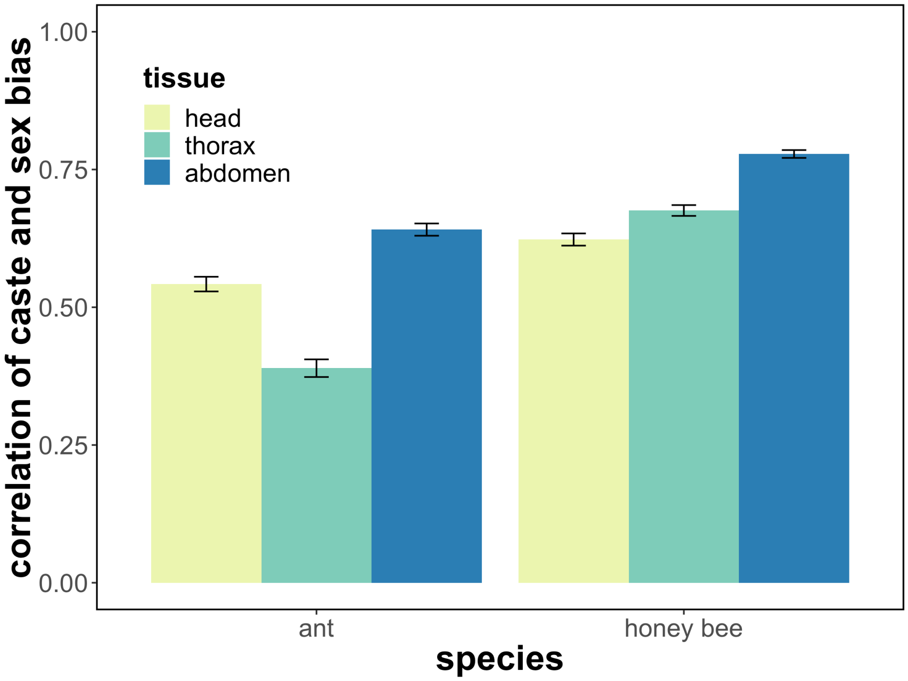


**Supplementary Figure 7. Caste-biased expression is correlated to sex-biased expression.** Pearson correlation of caste (queen/worker) and sex (queen/male) expression bias in ants and honey bees. Error bars represent Pearson correlation 95% confidence intervals. Correlations are significant in all cases (Pearson correlation; $P < 0.001$), but abdominal correlations are strongest. Source data are provided as a Source Data File, "Supplementary_Fig7.txt"

**Deleted: Supplementary Fig.**

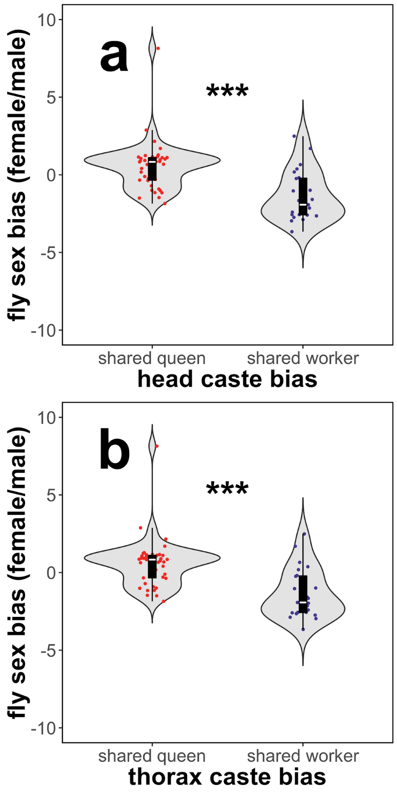


**Supplementary Figure 8. Social insect queen-biased genes tend to be female-biased in *D. melanogaster*.**

Deleted: Supplementary Fig.

Shared queen-biased DEGs tend to be female-biased in *D. melanogaster* while shared worker-biased DEGs tend to be male-biased in *D. melanogaster* (likely reflecting down-regulation in females) in both a) head and b) thoracic tissues. ). In each boxplot, the middle line represents median values, outer edges of boxplot represent upper and lower quartiles, and whiskers represent a deviation of 1.5*(interquartile range) from the upper and lower quartiles. Source data are provided as a Source Data File, "Supplementary_Fig8a.txt", "Supplementary_Fig8b.txt". *** = $P < 0.001$, Wilcoxon Test.

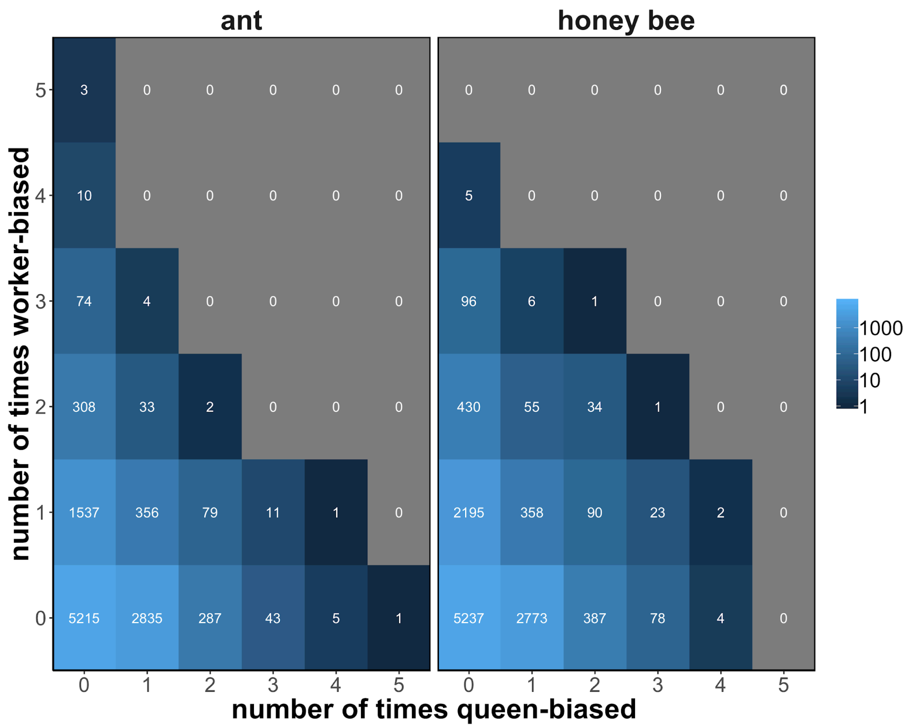


**Supplementary Figure 9. Distribution of the number of times genes exhibit biased expression towards each caste.**

Number of times each gene is upregulated in queen and workers across all comparisons (larva, pupa, and adult head, thorax, and abdomen). Color brightness is logarithmically proportional to the number of genes in each cell. N = 10804 genes for ants, N = 11775 genes for honey bees. Source data are provided as a Source Data File, "Supplementary_Fig9.txt".

Deleted: Supplementary Fig.

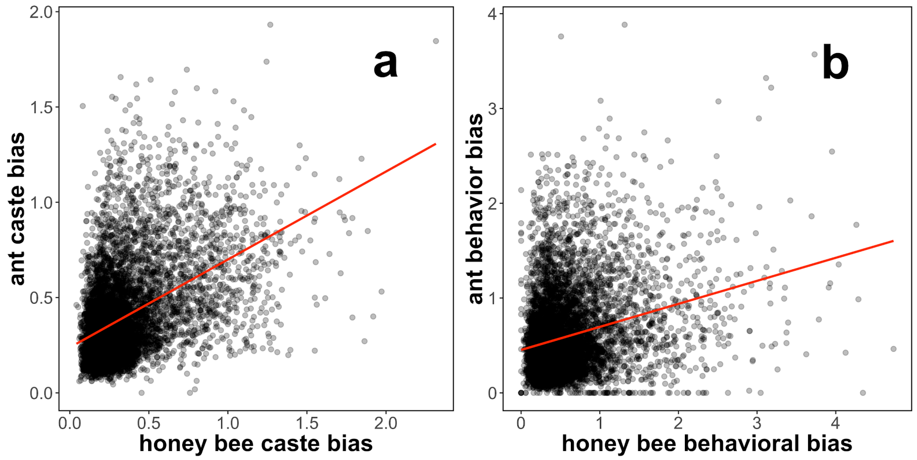


**Supplementary Figure 10. Caste and behavior bias are correlated between species.** Overall caste bias (a) and overall behavior bias (b) is correlated between ants and honey bees. “Overall” bias refers to the Euclidean distance of all $\log_2$ fold-change values (queens/worker for caste, nurses/foragers for behavior). The red line is the trendline of a linear model; Spearman correlation P < 0.001 in all cases. Source data are provided as a Source Data File, “Supplementary_Fig10a.txt”, “Supplementary_Fig10b.txt”.

**Deleted: Supplementary Fig.**

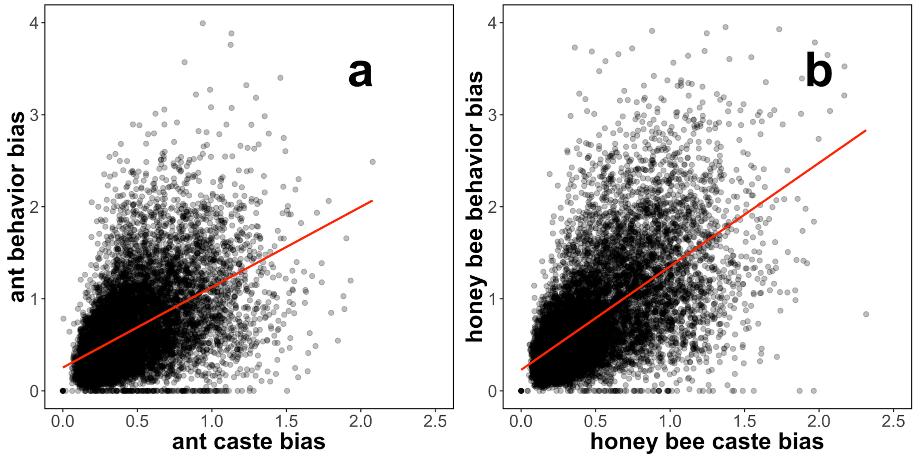


**Supplementary Figure 11. Caste bias is correlated to behavior bias.** Overall caste bias and overall behavior bias were correlated within a) ants and b) honey bees. "Overall" bias refers to the Euclidean distance of all $log_2$ fold-change values (queens/worker for caste, nurses/foragers for behavior). The red line is the trendline of a linear model; Spearman correlation $P < 0.001$ in all cases. Source data are provided as a Source Data File, "Supplementary_Fig11a.txt", "Supplementary_Fig11b.txt".

Deleted: Supplementary Fig.

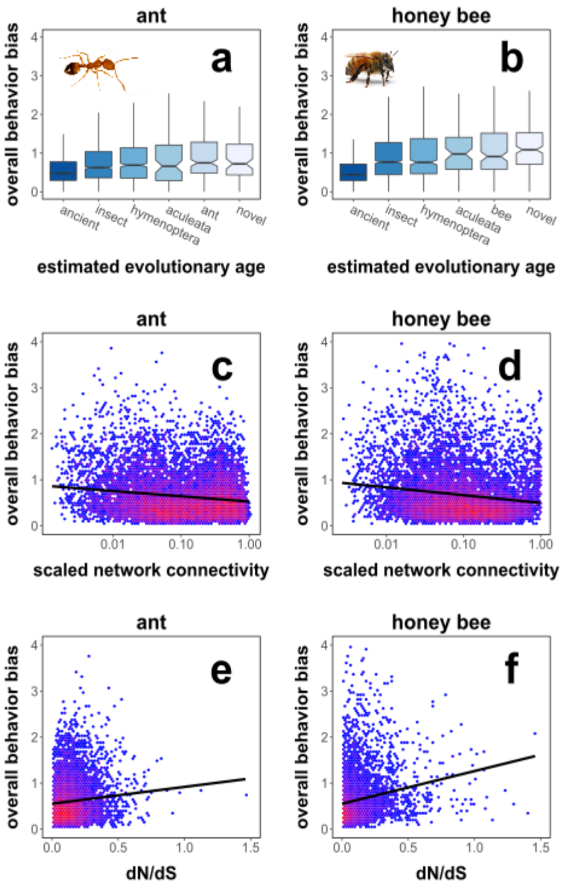


**Supplementary Figure 12. Evolutionary and network features of behavior-biased genes.** Genes that exhibit more behavior bias across tissues have younger estimated evolutionary ages (a,b) and tend to be loosely connected (c,d; Spearman correlation; ant: rho = -0.099, P < 0.001; honey bee: rho = -0.157, P < 0.001) and rapidly evolving (e,f; Spearman correlation; ant: rho = 0.079, P < 0.001; honey bee: rho = 0.226, P < 0.001). "Overall behavior bias" combines nurse forager $log_2$ fold-change values across all adult body segments. Connectivity is calculated using all samples and genes and scaled proportionally to the highest value. ). In each boxplot, the middle line represents median values, outer edges of boxplot represent upper and lower quartiles, and whiskers represent a deviation of 1.5*(interquartile range) from the upper and lower quartiles. Source data are provided as a Source Data File, "Supplementary_Fig12.txt". Photos were taken by Luigi Pontieri (pharaoh ant) and Alex Wild (honey bee).

Deleted: Supplementary Fig.

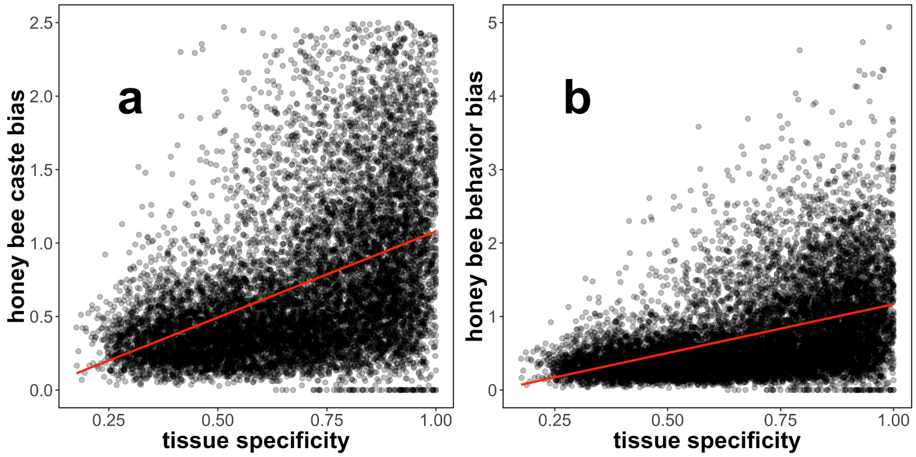


**Supplementary Figure 13. Caste and behavior bias are correlated to tissue specificity in honey bees.**

Deleted: Supplementary Fig.

Genes exhibiting more behavior bias tend to be tissue-specific. There was a positive correlation (Spearman correlation, $P < 0.001$ in each case) between caste/behavior bias and tissue specificity, where tissue specificity ($\tau$) is estimated using data from 12 honey bee tissues. $\tau = 1$ indicates a genes is expressed in only one tissue, while lower values indicate genes are more ubiquitously (i.e. evenly) expressed across tissues. The red line is the trendline of a linear model. Source data are provided as a Source Data File, "Supplementary_Fig13a.txt", "Supplementary_Fig13b.txt".

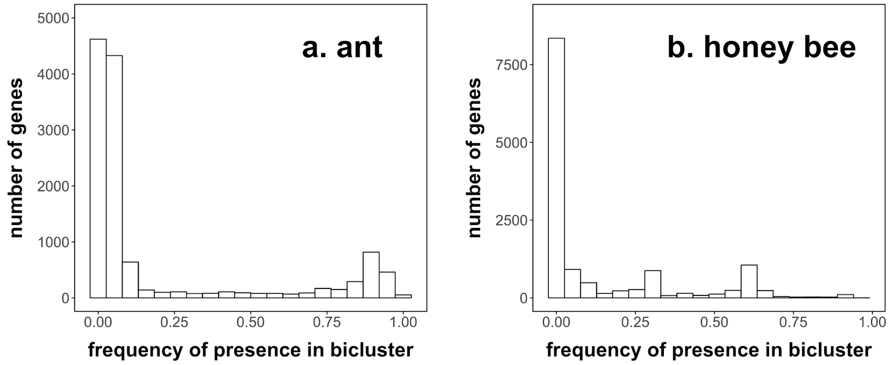


**Supplementary Figure 14. Distributions of the frequencies by which genes were placed in biclusters associated with queen abdomens.**

Histogram of the frequency with which genes were placed in the queen abdomen bicluster (out of 1000 runs). Plaid biclustering is a non-deterministic process, so different sets of genes can be present in each run. In ants (a), 1039 genes were present in >90% of queen abdomen biclusters and retained for further analysis. There are two peaks in the frequency distribution for honey bees (b). The lower frequency peak is made up of worker-associated genes (downregulated in queen abdomens) while the higher frequency peak (~60%) is made up of queen-associated genes. We retained genes with >60% frequency for further analysis. Source data are provided as a Source Data File, “Supplementary_Fig14a.txt”, “Supplementary_Fig14b.txt”.

Deleted: Supplementary Fig.

| Species | Stage | Tissue | Caste | Type | Replicates |
|---|---|---|---|---|---|
| ant | egg | whole body | N/A | N/A | 6 |
| ant | L1 | whole body | N/A | N/A | 6 |
| ant | L2 | whole body | queen | N/A | 3 |
| ant | L2 | whole body | worker | N/A | 3 |
| ant | L3 | whole body | queen | N/A | 3 |
| ant | L3 | whole body | worker | N/A | 3 |
| ant | L4 | whole body | queen | N/A | 3 |
| ant | L4 | whole body | worker | N/A | 3 |
| ant | L5 | whole body | queen | N/A | 3 |
| ant | L5 | whole body | worker | N/A | 3 |
| ant | pupa | whole body | queen | N/A | 3 |
| ant | pupa | whole body | worker | N/A | 3 |
| ant | pupa | whole body | male | N/A | 3 |
| ant | adult | head | queen | virgin queen | 3 |
| ant | adult | thorax | queen | virgin queen | 3 |
| ant | adult | abdomen | queen | virgin queen | 3 |
| ant | adult | head | queen | mated queen | 3 |
| ant | adult | thorax | queen | mated queen | 3 |
| ant | adult | abdomen | queen | mated queen | 3 |
| ant | adult | head | worker | nurse | 3 |
| ant | adult | thorax | worker | nurse | 3 |
| ant | adult | abdomen | worker | nurse | 3 |
| ant | adult | head | worker | forager | 3 |
| ant | adult | thorax | worker | forager | 3 |
| ant | adult | abdomen | worker | forager | 3 |
| ant | adult | head | male | N/A | 3 |
| ant | adult | thorax | male | N/A | 3 |
| ant | adult | abdomen | male | N/A | 3 |
| honey bee | egg | whole body | N/A | N/A | 3 |
| honey bee | L1 | whole body | N/A | N/A | 3 |
| honey bee | L2 | whole body | queen | N/A | 3 |
| honey bee | L2 | whole body | worker | N/A | 3 |
| honey bee | L3 | whole body | queen | N/A | 3 |
| honey bee | L3 | whole body | worker | N/A | 3 |
| honey bee | L4 | whole body | queen | N/A | 3 |
| honey bee | L4 | whole body | worker | N/A | 3 |
| honey bee | L5 | whole body | queen | N/A | 3 |
| honey bee | L5 | whole body | worker | N/A | 4 |
| honey bee | pupa | whole body | queen | N/A | 5 |
| honey bee | pupa | whole body | worker | N/A | 3 |
| honey bee | pupa | whole body | male | N/A | 3 |
| honey bee | adult | head | queen | virgin queen | 3 |
| honey bee | adult | thorax | queen | virgin queen | 3 |
| honey bee | adult | abdomen | queen | virgin queen | 3 |
| honey bee | adult | head | queen | mated queen | 3 |
| honey bee | adult | thorax | queen | mated queen | 3 |
| honey bee | adult | abdomen | queen | mated queen | 3 |
| honey bee | adult | head | worker | nurse | 3 |
| honey bee | adult | thorax | worker | nurse | 3 |
| honey bee | adult | abdomen | worker | nurse | 3 |
| honey bee | adult | head | worker | forager | 3 |
| honey bee | adult | thorax | worker | forager | 3 |
| honey bee | adult | abdomen | worker | forager | 3 |
| honey bee | adult | head | male | N/A | 3 |
| honey bee | adult | thorax | male | N/A | 3 |
| honey bee | adult | abdomen | male | N/A | 3 |

**Supplementary Table S1.** Full listing of sample types and number of each sample collected. "L1" and "L2" refers to larvae of the first and second stage, etc. We began caste-specific sampling at stage two because caste is determined and regulated in *M. pharaonis* by the end of the first larval instar[1]. After the first larval instar in *M. pharaonis*, worker-destined larvae can be distinguished from reproductive-destined larvae, which include male-destined and queen-destined larvae[1]. As such, our "queen-destined" ant larvae samples likely contain some male-destined larvae, but the proportion is expected to be low, as the sex ratio is known to be heavily queen-biased[2]. Sex and caste are both known in *A. mellifera* larvae, as individuals are reared in separate cells[3].

| stage/tissue | species | total DEGs | queen associated | worker associated |
|---|---|---|---|---|
| L2 | ant | 119 | 31 | 88 |
| L3 | ant | 818 | 490 | 328 |
| L4 | ant | 81 | 65 | 16 |
| L5 | ant | 757 | 437 | 320 |
| pupa | ant | 290 | 88 | 202 |
| head | ant | 741 | 420 | 321 |
| thorax | ant | 1327 | 695 | 632 |
| abdomen | ant | 4395 | 2711 | 1684 |
| larva_overall | ant | 361 | 241 | 120 |
| L2 | honey bee | 136 | 60 | 76 |
| L3 | honey bee | 117 | 28 | 89 |
| L4 | honey bee | 724 | 224 | 500 |
| L5 | honey bee | 1009 | 540 | 469 |
| pupa | honey bee | 245 | 163 | 82 |
| head | honey bee | 1144 | 717 | 427 |
| thorax | honey bee | 1369 | 721 | 648 |
| abdomen | honey bee | 5352 | 2769 | 2583 |
| larva_overall | honey bee | 473 | 176 | 297 |

**Supplementary Table S2.** Number of differentially expressed genes (DEGs) between queens and workers for each comparison (FDR < 0.1). "L2", "L3", etc refer to the 2nd and 3rd larval stage, respectively, while "larva_overall" is the result of differential expression with caste as main effect across all larval samples. Differentially expressed genes are divided into "queen associated", which exhibited higher expression in queens, and "worker associated", which exhibited higher expression in workers. Differential expression analysis performed with N = 10804 (ant) and 11775 (honey bee) genes.

| GO.ID | Term | P | stage/tissue |
|---|---|---|---|
| GO:0007265 | Ras protein signal transduction | 0.00026 | larva |
| GO:0046578 | regulation of Ras protein signal transduction | 0.00029 | larva |
| GO:0051056 | regulation of small GTPase mediated signal transduction | 0.00048 | larva |
| GO:0007264 | small GTPase mediated signal transduction | 0.00071 | larva |
| GO:0060628 | regulation of ER to Golgi vesicle-mediated transport | 0.00071 | larva |
| GO:0006970 | response to osmotic stress | 0.00089 | pupa |
| GO:0009651 | response to salt stress | 0.00287 | pupa |
| GO:0019432 | triglyceride biosynthetic process | 0.00297 | pupa |
| GO:0046460 | neutral lipid biosynthetic process | 0.00297 | pupa |
| GO:0046463 | acylglycerol biosynthetic process | 0.00297 | pupa |
| GO:0072525 | pyridine-containing compound biosynthetic process | 0.0018 | head |
| GO:0001704 | formation of primary germ layer | 0.0042 | head |
| GO:0002098 | tRNA wobble uridine modification | 0.0050 | head |
| GO:0043086 | negative regulation of catalytic activity | 0.0071 | head |
| GO:0010508 | positive regulation of autophagy | 0.0078 | head |
| GO:0019362 | pyridine nucleotide metabolic process | 0.00021 | thorax |
| GO:0046496 | nicotinamide nucleotide metabolic process | 0.00021 | thorax |
| GO:0006733 | oxidoreduction coenzyme metabolic process | 0.00043 | thorax |
| GO:0072524 | pyridine-containing compound metabolic process | 0.00054 | thorax |
| GO:0006739 | NADP metabolic process | 0.00088 | thorax |
| GO:0042445 | hormone metabolic process | 7.5e-05 | abdomen |
| GO:0010817 | regulation of hormone levels | 0.00041 | abdomen |
| GO:0042181 | ketone biosynthetic process | 0.00089 | abdomen |
| GO:0042180 | cellular ketone metabolic process | 0.00094 | abdomen |
| GO:0034754 | cellular hormone metabolic process | 0.00108 | abdomen |

**Supplementary Table S3.** Enriched gene ontology terms based on Gene Set Enrichment Analysis (GSEA) of differential expression between queens and workers in ants. P-value derived from Kolmogorov-Smirnov tests.

| GO.ID | Term | P | stage/tissue |
|---|---|---|---|
| GO:0019722 | calcium-mediated signaling | 0.0010 | larva |
| GO:0046113 | nucleobase catabolic process | 0.0021 | larva |
| GO:0007411 | axon guidance | 0.0024 | larva |
| GO:0061564 | axon development | 0.0027 | larva |
| GO:0035039 | male pronucleus assembly | 0.0032 | larva |
| GO:0008544 | epidermis development | 0.00053 | pupa |
| GO:0008286 | insulin receptor signaling pathway | 0.00065 | pupa |
| GO:0034599 | cellular response to oxidative stress | 0.00097 | pupa |
| GO:0032869 | cellular response to insulin stimulus | 0.00106 | pupa |
| GO:0071375 | cellular response to peptide hormone stimulus | 0.00106 | pupa |
| GO:0008610 | lipid biosynthetic process | 0.00011 | head |
| GO:0016070 | RNA metabolic process | 0.00018 | head |
| GO:0030534 | adult behavior | 0.00028 | head |
| GO:0008344 | adult locomotory behavior | 0.00031 | head |
| GO:0007478 | leg disc morphogenesis | 0.00041 | head |
| GO:0048747 | muscle fiber development | 0.0011 | thorax |
| GO:0090254 | cell elongation involved in imaginal disc-derived wing morphogenesis | 0.0027 | thorax |
| GO:0006457 | protein folding | 0.0045 | thorax |
| GO:0071897 | DNA biosynthetic process | 0.0055 | thorax |
| GO:0034063 | stress granule assembly | 0.0071 | thorax |
| GO:0045887 | positive regulation of synaptic growth at neuromuscular junction | 0.00037 | abdomen |
| GO:1904398 | positive regulation of neuromuscular junction development | 0.00037 | abdomen |
| GO:0051965 | positive regulation of synapse assembly | 0.00116 | abdomen |
| GO:0030490 | maturation of SSU-rRNA | 0.00126 | abdomen |
| GO:0007436 | larval salivary gland morphogenesis | 0.00205 | abdomen |

**Supplementary Table S4.** Enriched gene ontology terms based on Gene Set Enrichment Analysis (GSEA) of differential expression between queens and workers in honey bees. P-value derived from Kolmogorov-Smirnov tests.

| stage/tissue | species | total DEGs | nurse associated | forager associated |
|---|---|---|---|---|
| head | ant | 405 | 314 | 91 |
| thorax | ant | 490 | 305 | 185 |
| abdomen | ant | 544 | 341 | 203 |
| head | honey bee | 927 | 404 | 523 |
| thorax | honey bee | 2519 | 1243 | 1276 |
| abdomen | honey bee | 2017 | 1007 | 1010 |

**Supplementary Table S5.** Number of differentially expressed genes (DEGs) between nurses and foragers for each comparison (FDR < 0.1). Differentially expressed genes are divided into "nurse associated", which exhibited higher expression in nurses, and "forager associated", which exhibited higher expression in foragers. Differential expression analysis performed with N = 10804 (ant) and 11775 (honey bee) genes.

| GO.ID | Term | P | tissue |
|---|---|---|---|
| GO:0048284 | organelle fusion | 0.00021 | head |
| GO:0006629 | lipid metabolic process | 0.00138 | head |
| GO:0048580 | regulation of post-embryonic development | 0.00186 | head |
| GO:0044255 | cellular lipid metabolic process | 0.00217 | head |
| GO:0044801 | single-organism membrane fusion | 0.00226 | head |
| GO:0032502 | developmental process | 0.00019 | thorax |
| GO:0007525 | somatic muscle development | 0.00035 | thorax |
| GO:0044767 | single-organism developmental process | 0.00041 | thorax |
| GO:0007275 | multicellular organism development | 0.00041 | thorax |
| GO:0090175 | regulation of establishment of planar polarity | 0.00043 | thorax |
| GO:0000289 | nuclear-transcribed mRNA poly(A) tail shortening | 0.0017 | abdomen |
| GO:0007006 | mitochondrial membrane organization | 0.0054 | abdomen |
| GO:0042441 | eye pigment metabolic process | 0.0056 | abdomen |
| GO:0043324 | pigment metabolic process involved in developmental pigmentation | 0.0056 | abdomen |
| GO:0043474 | pigment metabolic process involved in pigmentation | 0.0056 | abdomen |

**Supplementary Table S6.** Enriched gene ontology terms based on Gene Set Enrichment Analysis (GSEA) of differential expression between nurses and foragers in ants. P-value derived from Kolmogorov-Smirnov tests.

| GO.ID | Term | P | stage/tissue |
|---|---|---|---|
| GO:0019722 | calcium-mediated signaling | 0.0010 | larva |
| GO:0046113 | nucleobase catabolic process | 0.0021 | larva |
| GO:0007411 | axon guidance | 0.0024 | larva |
| GO:0061564 | axon development | 0.0027 | larva |
| GO:0035039 | male pronucleus assembly | 0.0032 | larva |
| GO:0008544 | epidermis development | 0.00053 | pupa |
| GO:0008286 | insulin receptor signaling pathway | 0.00065 | pupa |
| GO:0034599 | cellular response to oxidative stress | 0.00097 | pupa |
| GO:0032869 | cellular response to insulin stimulus | 0.00106 | pupa |
| GO:0071375 | cellular response to peptide hormone stimulus | 0.00106 | pupa |
| GO:0008610 | lipid biosynthetic process | 0.00011 | head |
| GO:0016070 | RNA metabolic process | 0.00018 | head |
| GO:0030534 | adult behavior | 0.00028 | head |
| GO:0008344 | adult locomotory behavior | 0.00031 | head |
| GO:0007478 | leg disc morphogenesis | 0.00041 | head |
| GO:0048747 | muscle fiber development | 0.0011 | thorax |
| GO:0090254 | cell elongation involved in imaginal disc-derived wing morphogenesis | 0.0027 | thorax |
| GO:0006457 | protein folding | 0.0045 | thorax |
| GO:0071897 | DNA biosynthetic process | 0.0055 | thorax |
| GO:0034063 | stress granule assembly | 0.0071 | thorax |
| GO:0045887 | positive regulation of synaptic growth at neuromuscular junction | 0.00037 | abdomen |
| GO:1904398 | positive regulation of neuromuscular junction development | 0.00037 | abdomen |
| GO:0051965 | positive regulation of synapse assembly | 0.00116 | abdomen |
| GO:0030490 | maturation of SSU-rRNA | 0.00126 | abdomen |
| GO:0007436 | larval salivary gland morphogenesis | 0.00205 | abdomen |

**Supplementary Table S7.** Enriched gene ontology terms based on Gene Set Enrichment Analysis (GSEA) of differential expression between nurses and foragers in honey bees. P-value derived from Kolmogorov-Smirnov tests.

| Gene | logFC queen/worker | connectivity | SwissProt |
|---|---|---|---|
| LOC105837185 | 9.333 | 0.793 | Leukocyte elastase inhibitor A |
| LOC105838268 | 8.610 | 0.790 | Gephyrin |
| LOC105836111 | 8.147 | 0.784 | RCC1 and BTB domain-containing protein 1 |
| LOC105836023 | 7.962 | 0.772 | Histone H2B |
| LOC105834654 | 7.258 | 0.797 | Vitellogenin receptor |
| LOC105837528 | 6.871 | 0.848 | Ankyrin-2 |
| LOC105838623 | 6.713 | 0.855 | Transcription factor SOX-14 |
| LOC105829700 | 6.447 | 0.839 | Maternal embryonic leucine zipper kinase |
| LOC105834441 | 6.307 | 0.864 | Nuclear RNA export factor 1 |
| LOC105830728 | 6.115 | 0.911 | S-phase kinase-associated protein 2 |
| LOC105837219 | 5.825 | 0.926 | Rac GTPase-activating protein 1 |
| LOC105840891 | 5.777 | 0.973 | Acidic repeat-containing protein |
| LOC105838786 | 5.389 | 0.824 | Spondin-1 |
| LOC105837988 | 5.364 | 0.819 | Multiple PDZ domain protein |
| LOC105830806 | 5.311 | 0.833 | Insulin-degrading enzyme |
| LOC105836312 | 5.144 | 0.797 | Serine protease nudel |
| LOC105836129 | 5.108 | 0.915 | Rho GTPase-activating protein 19 |
| LOC105834586 | 5.078 | 0.906 | Putative bifunctional UDP-N-acetylglucosamine transferase and deubiquitinase ALG13 |
| LOC105832223 | 5.058 | 0.874 | E3 ubiquitin-protein ligase SIAH1 |
| LOC105840292 | 4.974 | 0.785 | Pre-mRNA-splicing factor RBM22 |
| LOC105840093 | 4.833 | 0.897 | ATP-dependent RNA helicase vasa isoform A |
| LOC105833898 | 4.812 | 0.925 | Piwi-like protein 1 |
| LOC105839662 | 4.676 | 0.966 | G2/mitotic-specific cyclin-B3 |
| LOC105828383 | 4.542 | 0.949 | Histone RNA hairpin-binding protein |
| LOC105831777 | 4.326 | 0.815 | Protein dispatched |
| LOC105835848 | 4.292 | 0.807 | Glyoxylate reductase |
| LOC105838831 | 4.242 | 0.890 | Broad-complex core protein isoform 6 |
| LOC105832464 | 4.188 | 0.774 | Zinc finger protein 800 |
| LOC105837226 | 4.096 | 0.883 | DNA repair and recombination protein RAD54-like (Fragment) |
| LOC105834656 | 4.077 | 0.805 | Putative ATP-dependent RNA helicase me31b |

**Supplementary Table S8.** Hub genes of the queen abdominal module in ants. Hub genes were defined as genes with intra-modular connectivity in at least the 90th percentile, and $\log_2$ fold-change (queen/worker) greater than 2.

| Gene | logFC queen/worker | connectivity | SwissProt |
|---|---|---|---|
| LOC724752 | 11.662 | 0.746 | E3 ubiquitin-protein ligase TRIM71 |
| LOC410888 | 9.000 | 0.829 | lachesin-like |
| LOC410684 | 7.405 | 0.754 | homeobox protein OTX1 A |
| LOC100576333 | 6.493 | 0.738 | nucleoredoxin-like |
| LOC725841 | 6.339 | 0.897 | hyaluronan mediated motility receptor |
| LOC100577382 | 6.272 | 0.896 | targeting protein for Xklp2 homolog |
| LOC551099 | 6.172 | 0.760 | coiled-coil domain-containing protein 43 |
| LOC724193 | 5.957 | 0.859 | kinesin-like protein KIF18A |
| LOC726506 | 5.927 | 0.828 | protein claret segregational |
| LOC725920 | 5.829 | 0.875 | vitellogenin receptor |
| LOC100576828 | 5.826 | 0.843 | protein maelstrom 2 |
| LOC412031 | 5.811 | 0.930 | S-phase kinase-associated protein 2 |
| LOC102656846 | 5.705 | 0.942 | cyclin-A2 |
| LOC411529 | 5.531 | 0.861 | maternal embryonic leucine zipper kinase-like |
| LOC410502 | 5.456 | 0.910 | transformation/transcription domain-associated protein |
| LOC410015 | 5.270 | 0.778 | protein LSM14 homolog A |
| LOC100578691 | 5.186 | 0.909 | rhoGEF domain-containing protein gxcJ-like |
| LOC100578255 | 5.073 | 0.734 | ras GTPase-activating-like protein IQGAP1 |
| LOC552100 | 4.953 | 0.809 | protein ovo |
| LOC100576908 | 4.883 | 0.917 | polycomb protein Asx |
| LOC551871 | 4.808 | 0.912 | P protein-like |
| LOC411970 | 4.546 | 0.767 | G kinase-anchoring protein 1-like |
| LOC725606 | 4.498 | 0.758 | serine protease gd |
| LOC409092 | 4.486 | 0.850 | enhancer of mRNA-decapping protein 3 |
| LOC409681 | 4.483 | 0.807 | RWD domain-containing protein 1 |
| LOC411809 | 4.460 | 0.780 | enolase-phosphatase E1 |
| LOC551773 | 4.320 | 0.876 | serine/threonine-protein kinase VRK1-like |
| LOC413667 | 4.212 | 0.914 | G2/mitotic-specific cyclin-B3 |
| LOC409472 | 3.995 | 0.975 | protein Smaug homolog 1 |
| LOC552725 | 3.868 | 0.848 | N-acetylglucosamine-1-phosphotransferase subunits alpha/beta |

**Supplementary Table S9.** Hub genes of the queen abdominal module in honey bees. Hub genes were defined as genes with intra-modular connectivity in at least the 90th percentile, and $\log_2$ fold-change (queen/worker) greater than 2.

| GO.ID | Term | P | test |
|---|---|---|---|
| GO:0010977 | negative regulation of neuron projection development | 0.00032 | ant caste |
| GO:0014017 | neuroblast fate commitment | 0.00038 | ant caste |
| GO:0045165 | cell fate commitment | 0.00041 | ant caste |
| GO:0007400 | neuroblast fate determination | 0.00085 | ant caste |
| GO:0010771 | negative regulation of cell morphogenesis involved in differentiation | 0.00122 | ant caste |
| GO:0060322 | head development | 2.3e-07 | ant behavior |
| GO:0007420 | brain development | 3.6e-07 | ant behavior |
| GO:0016319 | mushroom body development | 1.6e-05 | ant behavior |
| GO:0045165 | cell fate commitment | 2.1e-05 | ant behavior |
| GO:0007417 | central nervous system development | 4.2e-05 | ant behavior |
| GO:0006650 | glycerophospholipid metabolic process | 0.00046 | bee caste |
| GO:0051231 | spindle elongation | 0.00098 | bee caste |
| GO:0046488 | phosphatidylinositol metabolic process | 0.00134 | bee caste |
| GO:0035050 | embryonic heart tube development | 0.00194 | bee caste |
| GO:0000022 | mitotic spindle elongation | 0.00222 | bee caste |
| GO:0035295 | tube development | 0.00020 | bee behavior |
| GO:0002009 | morphogenesis of an epithelium | 0.00029 | bee behavior |
| GO:0035239 | tube morphogenesis | 0.00032 | bee behavior |
| GO:0048729 | tissue morphogenesis | 0.00061 | bee behavior |
| GO:0060562 | epithelial tube morphogenesis | 0.00123 | bee behavior |

**Supplementary Table S10.** Enriched gene ontology terms based on overall caste or behavior bias in ants and honey bees. GO terms are derived from *D. melanogaster* orthologs. P-value is from gene set enrichment analysis (Kolmogorov–Smirnov test)

| species | comparison | abdomen included? | variable tested | Spearman rho | P-value |
|---|---|---|---|---|---|
| ant | caste | yes | connectivity | -0.162 | 3.92e-33 |
| ant | caste | no | connectivity | -0.282 | 1.38e-99 |
| ant | caste | yes | dN/dS | 0.151 | 6.07e-29 |
| ant | caste | no | dN/dS | 0.130 | 8.51e-22 |
| ant | caste | yes | evolutionary age | 0.159 | 5.19e-32 |
| ant | caste | no | evolutionary age | 0.161 | 5.96e-33 |
| honey bee | caste | yes | connectivity | 0.003 | 8.13e-01 |
| honey bee | caste | no | connectivity | -0.132 | 4.61e-20 |
| honey bee | caste | yes | dN/dS | 0.155 | 3.15e-27 |
| honey bee | caste | no | dN/dS | 0.169 | 4.27e-32 |
| honey bee | caste | yes | evolutionary age | 0.166 | 3.72e-31 |
| honey bee | caste | no | evolutionary age | 0.154 | 7.95e-27 |
| honey bee | caste | yes | tau | 0.409 | 1.33e-193 |
| honey bee | caste | no | tau | 0.397 | 1.14e-180 |
| ant | behavior | yes | connectivity | -0.157 | 4.29e-31 |
| ant | behavior | no | connectivity | -0.119 | 1.24e-18 |
| ant | behavior | yes | dN/dS | 0.043 | 1.75e-03 |
| ant | behavior | no | dN/dS | 0.020 | 1.38e-01 |
| ant | behavior | yes | evolutionary age | 0.021 | 1.25e-01 |
| ant | behavior | no | evolutionary age | 0.014 | 3.02e-01 |
| honey bee | behavior | yes | connectivity | -0.162 | 9.22e-30 |
| honey bee | behavior | no | connectivity | -0.137 | 1.49e-21 |
| honey bee | behavior | yes | dN/dS | 0.174 | 7.02e-34 |
| honey bee | behavior | no | dN/dS | 0.150 | 1.83e-25 |
| honey bee | behavior | yes | evolutionary age | 0.161 | 3.49e-29 |
| honey bee | behavior | no | evolutionary age | 0.126 | 1.95e-18 |
| honey bee | behavior | yes | tau | 0.328 | 9.55e-121 |
| honey bee | behavior | no | tau | 0.281 | 5.79e-88 |

**Supplementary Table S11.** Partial correlation between connectivity, evolutionary rate (dN/dS), evolutionary age (phylostrata), and tissue-specificity (tau) and caste or behavior bias while accounting for expression. Analysis was performed separately for each species and comparison (i.e. separately for caste bias and behavior bias), as well as while including or excluding abdomen in calculations of caste bias and expression. Connectivity is total connectivity measured across all samples and genes. Phylostrata is a measure of estimated evolutionary age, with higher values indicating younger genes. Tau is the degree to which genes exhibit tissue-specific expression across 12 honey bee tissues (results presented only for honey bees). N = 10520 genes (ants), 10011 genes (honey bees). To estimate a measure of expression analogous to overall bias, we calculated the Euclidean distance of $\log_{10}$ counts-per-million at each stage/tissue tested.

**Supplementary Table S12.** List of species used for phylostratigraphy analysis, with the NCBI Taxonomy ID.

| Species | NCBI Taxonomy ID |
|---|---|
| Acromyrmex echinatior | 103372 |
| Atta cephalotes | 12957 |
| Atta colombica | 520822 |
| Camponotus floridanus | 104421 |
| Cardiocondyla obscurior | 286306 |
| Monomorium pharaonis | 307658 |
| Linepithema humile | 83485 |
| Lasius niger | 67767 |
| Harpegnathos saltator | 610380 |
| Dinoponera quadriceps | 609295 |
| Cyphomyrmex costatus | 456900 |
| Ooceraea biroi | 2015173 |
| Pogonomyrmex barbatus | 144034 |
| Pseudomyrmex gracilis | 219809 |
| Solenopsis invicta | 13686 |
| Trachymyrmex septentrionalis | 34720 |
| Trachymyrmex cornetzi | 471704 |
| Trachymyrmex zeteki | 64791 |
| Vollenhovia emeryi | 411798 |
| Wasmannia auropunctata | 64793 |
| Temnothorax curvispinosus | 300111 |
| Apis cerana | 7461 |
| Apis dorsata | 7462 |
| Apis florea | 7463 |
| Apis mellifera | 7460 |
| Bombus impatiens | 132113 |
| Bombus terrestris | 30195 |
| Melipona quadrifasciata | 166423 |
| Eufriesea mexicana | 516756 |
| Ceratina calcarata | 156304 |
| Megachile rotundata | 143995 |
| Habropoda laboriosa | 597456 |
| Dufourea novaeangliae | 178035 |
| Polistes canadensis | 91411 |
| Polistes dominula | 743375 |
| Ceratosolen solmsi | 142686 |
| Ceratosolen solmsi marchali | 326594 |
| Copidosoma floridanum | 29053 |
| Fopius arisanus | 64838 |
| Microplitis demolitor | 69319 |
| Nasonia vitripennis | 7425 |
| Trichogramma pretiosum | 7493 |
| Trichomalopsis sarcophagae | 543379 |
| Diachasma alloeum | 454923 |
| Orussus abietinus | 222816 |
| Athalia rosae | 37344 |
| Cephus cinctus | 211228 |
| Neodiprion lecontei | 441921 |
| Pediculus humanus | 121225 |
| Drosophila melanogaster | 7227 |
| Aedes aegypti | 7159 |
| Bombyx mori | 7091 |
| Papilio machaon | 76193 |
| Anopheles gambiae | 7165 |
| Onthophagus taurus | 166361 |
| Tribolium castaneum | 7070 |
| Acyrthosiphon pisum | 7029 |
| Zootermopsis nevadensis | 136037 |
| Caenorhabditis elegans | 6239 |
| Hydra vulgaris | 6087 |
| Strongylocentrotus purpuratus | 7668 |
| Lottia gigantea | 225164 |
| Helobdella robusta | 6412 |
| Mus musculus | 10090 |
| Homo sapiens | 9606 |
| Xenopus tropicalis | 8364 |
| Latimeria chalumnae | 7897 |
| Danio rerio | 7955 |
